# Potato virus X -delivered CRISPR activation programs lead to strong endogenous gene induction and transient metabolic reprogramming in *Nicotiana benthamiana*

**DOI:** 10.1101/2022.04.21.489058

**Authors:** S Selma, S Gianoglio, M Uranga, M Vázquez-Vilar, A Espinosa-Ruiz, M Drapal, PD Fraser, JA Daròs, D Orzaez

## Abstract

Programmable transcriptional regulators based on CRISPR architecture are promising tools for the control of plant gene expression. In plants, CRISPR gene activation (CRISPRa) has been shown effective in modulating development processes, such as the flowering time, or customising biochemical composition. The most widely used method for delivering the CRISPR components into the plant is *Agrobacterium tumefaciens*-mediated genetic transformation, either transient or stable. However, due to their versatility and their ability to move, virus-derived systems have emerged as an interesting alternative for supplying the CRISPR components to the plant, in particular the gRNA, which represents the variable component in CRISPR strategies. In this work we describe a *Potato virus X* (PVX)-derived vector that, upon agroinfection in *N. benthamiana*, serves as a vehicle for gRNAs delivery, producing a highly specific Virus-Induced Gene Activation (VIGA). The system works in combination with a *Nicotiana benthamiana* transgenic line carrying the remaining complementary CRISPRa components, specifically the dCasEV2.1 cassette, which has previously been shown to mediate strong programmable transcriptional activation in plants. Using an easily scalable, non-invasive spraying method, we show here that gRNAs-mediated activation programs move locally and systemically generating a strong activation response in different target genes. Furthermore, by activating three different endogenous MYB transcription factors, we demonstrate that this PVX-based virus-induced gene reprogramming (VIGR) strategy results in program-specific metabolic fingerprints in *N. benthamiana* leaves characterized by distinctive phenylpropanoid-enriched metabolite profiles.

## Introduction

The emerging CRISPR-Cas systems provide a battery of specific and efficient tools for gene editing and regulation (Mao et al. 2019; Moradpour et al. 2020). The recent achievements in targeted gene regulation in plants employing CRISPR/Cas-based programmable transcription factors not only enhance our ability to accurately explore the links between gene expression and phenotype (Lowder et al. 2018; Papikian et al. 2019; Pan et al. 2021b) but also open a new paradigm for targeted crop improvement by, for example, tuning flowering time or customizing metabolic composition (Lessard et al. 2002; Charfeddine et al. 2019; Maeda and Nakamichi 2022). CRISPR-based transcriptional regulators typically comprise a catalytically inactive Cas enzyme with transcriptional regulatory domains anchored to its structure. This allows the specific binding of the chimeric transcriptional regulator to the genomic DNA, guided by the 20 nucleotides of the guide RNAs (gRNAs) (Lee et al. 2019). The most widely used Cas nuclease is the Cas9 endonuclease from *Streptococcus pyogenes*, however other nucleases have been also optimized for achieving transcriptional regulation in plants (Zhang et al. 2021c). Currently, engineering efforts are focused on improving the efficiency and specificity of programmed gene regulation and also on the development of optimized delivery methods for the information encoded in the gRNA sequence. Thus, in an ideal scenario, crop plants could contain integrated CRISPR systems capable of executing gene activation instructions that would be delivered in the form of gRNAs commands that move systemically throughout the plant (Molina-Hidalgo et al. 2020).

Initial approaches for the delivery of CRISPR components to the plant were based on the widely used *Agrobacterium*-mediated T-DNA transformation, biolistic-delivery or protoplast transformation methods (Mathur and Koncz 1998; Wu et al. 2020). The *Agrobacterium*-mediated transformation relies on already established plant transformation protocols (Tzfira and Citovsky 2006; Danilo et al. 2019). It is easy to handle and does not present strict limitations on the size of DNA constructs, but it requires the generation of stable transgenics through tissue culture and regeneration. *Agrobacterium*-mediated transformation can operate transiently in some species, such as *N. benthamiana* (Wydro et al. 2006), without generating stable transgenic lines. In this case, however, the action of the CRISPR elements will be limited to the area of agroinfiltration. The biolistic delivery and protoplast transfection methods provide alternatives for plant species unable to be transformed with *Agrobacterium*, and offer non-transgenic options for genome editing (Hamada et al. 2017), like the preassembling of the Cas protein and the gRNA to form ribonucleoproteins (RNPs) (Liang et al. 2019; Zhang et al. 2021b). However, these delivery strategies present some disadvantages, such as the instability and the rapid degradation of the RNP complex, and do not circumvent the requirements of plant regeneration, which is not suitable for all plant species.

Classic work showed that viruses can be used to silence plant endogenous genes by simply harbouring a sequence fragment homologous to the target gene in an approaches known as virus induced-gene silencing (VIGS) (Lu et al. 2003). In addition, recent studies highlight the potential use of viral vectors as transient delivery vehicles for CRISPR-Cas components in many biological systems including plants (Platt et al. 2014; Senís et al. 2014; Xu et al. 2019). This approach, commonly termed virus-induced genome editing (VIGE), has mainly focused on the delivery of one or more gRNAs using RNA or DNA virus vectors in transgenic plants that stably express the Cas nuclease (Gentzel et al. 2022). The range of viral systems used for gRNA-delivery in VIGE approaches is expanding, also enlarging the range of suitable plant hosts. Recently described VIGE vectors include the *Tobacco rattle virus* (TRV) (Ali et al. 2015; Ellison et al. 2020), the *Cabbage Leaf Curl virus* (CaLCuV) (Yin et al. 2015), the *Tobacco mosaic virus* (TMV) (Cody et al. 2017), the *Pea early browning virus* (Ali et al. 2018), the *Foxtail mosaic virus* (FoMV) (Mei et al. 2019), the *Barley stripe mosaic virus* (BSMV) (Hu et al. 2019; Li et al. 2021) and the *Beet necrotic yellow vein virus* (BNYVV) (Jiang et al. 2020). One of the previously reported VIGE systems that showed remarkable efficiency in terms of gene editing was based on the *Potato virus X* (PVX) (Ariga et al. 2020), a member of the genus Potexvirus that infects 62 plant species, several of them belonging to the Solanaceae family (Lico et al. 2015). In addition, the PVX infection can be easily traced due to the phenotypic alterations produced in the host in a relatively short period. This recombinant virus was engineered to drive the expression of the gRNA by the sub-genomic coat protein (CP) promoter. Also, this viral system entails a multiplexing strategy for gRNA expression with the particularity that the PVX-based gRNA delivery does not need the presence of a tRNA-processing system for properly expressing several gRNAs in tandem (Uranga et al. 2021a).

Viral vectors have been also proposed as shuttles for the transient delivery of exogenous, RNA-encoded information to plant crops, leading to the ectopic and generalized expression of, for example, added-value recombinant proteins, defence-related genes or developmental regulators (Marillonnet et al. 2005; Gleba et al. 2013; Torti et al. 2021). This inspiring new breeding concept of transient genetic reprogramming can be expanded even further with the introduction of CRISPR programmable regulators. In this newly proposed scheme, gRNAs delivered to the plant using viral vectors would serve to reprogram endogenous gene expression. In the present study, we aimed to explore the potential of PVX -based gRNA delivery for strong customized transcriptional activation in plants, thus expanding the toolbox of viral systems employed for gene regulation, which is currently based only on TRV (Ghoshal et al. 2020; Khakhar et al. 2021). To achieve this, the PVX-based VIGE vector developed by Uranga et al. (2021) was further engineered to express tandem repeats of modified gRNAs incorporating two RNA aptamers that bind the coat protein of the MS2 phage. This modification is an adaptation to the so-called dCasEV2.1 system, a strong CRISPR programable activator earlier described by our group (Selma et al. 2019). In addition, the constant elements of the dCasEV2.1 system were stably transformed in *N. benthamiana*. The constant dCasEV2.1 module comprises two transcriptional units (TUs) driven by the *Cauliflower mosaic virus* (CaMV) 35S promoter: (i) a TU encoding an inactive version of the Cas9 (dCas9) fused to the plant activation domain EDLL (Tiwari et al. 2012), and (ii) a second TU encoding the coat protein of the MS2 phage fused to the VPR activator. VPR is a tandem fusion domain comprising the viral transactivation domains VP64, P65 and Rta (Chavez et al. 2015). Through a spray-based agrodelivery method that is easy to handle, non-invasive, and adapted for large scale applications, we demonstrated that the modified gRNAs were efficiently delivered to the plant, leading to a potent PVX-based virus-induced gene reprogramming (VIGR). By targeting three different endogenous MYB factors (*NbODO1, NbMYB24, and NbMYB21*), allegedly involved in regulating the phenylpropanoid pathway, we generated three different phenylpropanoid-enriched chemotypes in *N. benthamiana* leaves, obtaining information on the specific transcriptomic and metabolic fingerprint produced by each MYB factor.

## Results

### Generation of dCasEV2.1 *N. benthamiana* chassis for CRISPRa

The reported ability of CRISPR-based transcriptional activators to efficiently regulate gene expression in *N. benthamiana* prompted us to generate a transgenic line to serve as the chassis for CRISPR-mediated targeted transcriptional activation (CRISPRa) experiments. To achieve this, a construct carrying the constant genetic elements in the dCasEV2.1 strategy, along with a yellow fluorescent protein (YFP) (Figure 1A), was employed to generate the new transgenic lines, named YFP-dCasEV. Upon transient delivery to the plant cells of one or more gRNAs targeting a specific promoter, a full dCasEV2.1 ribonucleoprotein complex will be assembled, which would expectedly result in a strong and specific gene activation (Figure 1B). The YFP-positive transgenic plants obtained in Agrobacterium-mediated transformation were then evaluated in a CRISPRa assay where a gRNA targeting the tomato *Dihydroflavonol 4-reductase* (*SlDFR*) promoter was delivered via agroinfiltration. Targeted activation was scored using a luciferase reporter driven by the *SlDFR* promoter and co-delivered together with the gRNA. Six transgenic lines were generated, but only two of them, namely YFP-dCasEV_L3 and YFP-dCasEV_L6, showed a strong gene activation upon specific gRNA expression and no activation in the absence of gRNA. As a positive control, a wild type *N. benthamiana* line was transiently co-infiltrated with dCas9EV2.1 module, gRNA, and the luciferase reporter. No statistically significant differences were found between the two positives, stably transformed lines and the transiently infiltrated control (Figure 1C). We decided to establish YFP-dCasEV_L6 as our chassis line, from which we later obtained a homozygous T2 population.

**Figure 1:**
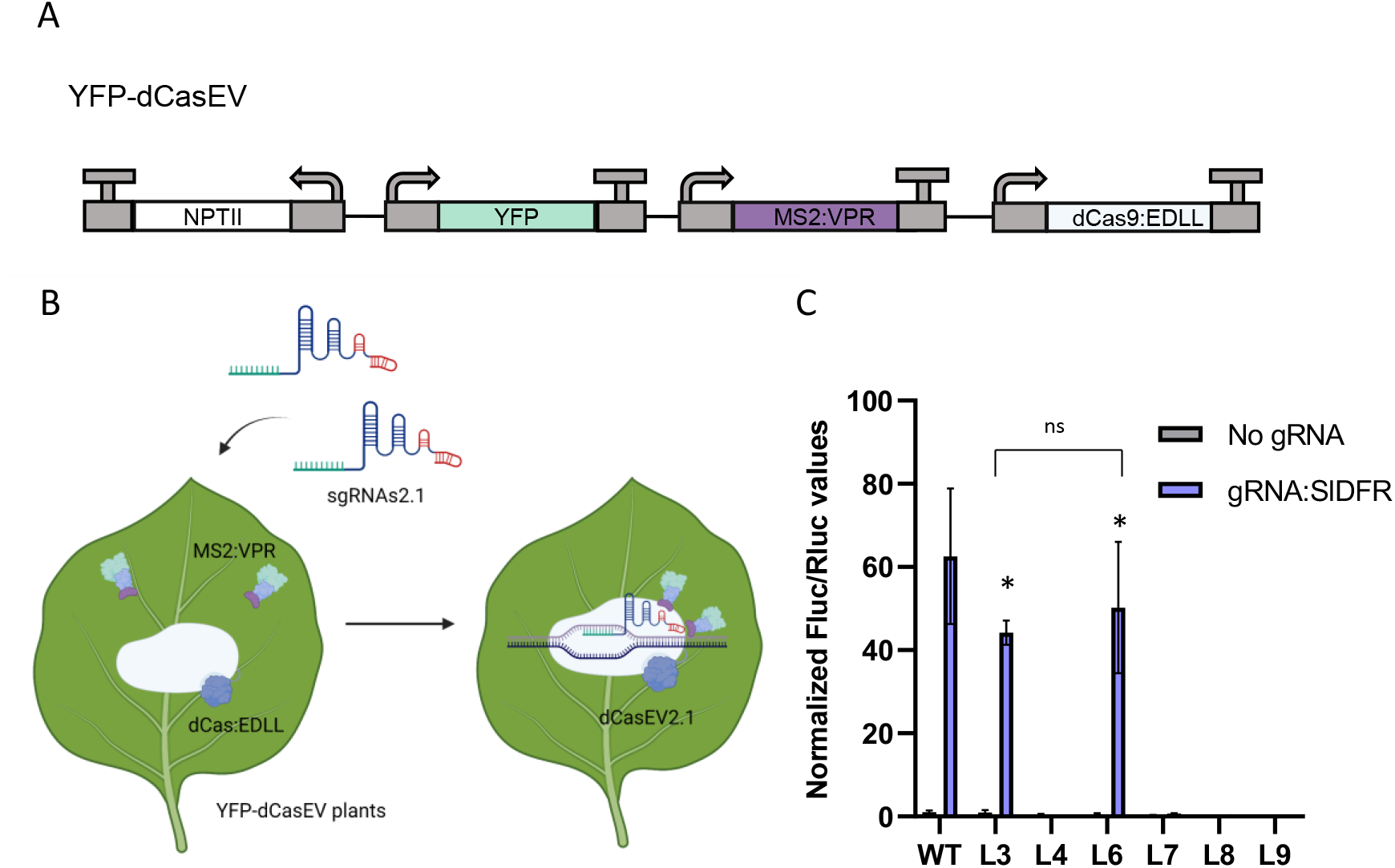
Generation and selection of *N. benthamiana* chassis for CRISPRa. A) Representation of the DNA construct employed for *N. benthamiana* transformation that includes the constant module of the dCasEV2.1 (dCas9:EDLL and MS2:VPR), a YFP reporter (YFP), and a Kanamycin resistance (*NPTII*) gene. B) Schematic representation of action mechanism of YFP-dCasEV plants by the exogenous incorporation of gRNAs (gRNA2.1), forming the dCasEV2.1 complex as a transcriptional activation system. C) Normalized Fluc/ Rluc ratios measured in transgenic lines containing the constant module of dCasEV2.1. The right side of the leaf transiently expresses a luciferase reporter driven by the *SlDFR* promoter and a gRNA that targets the position -150 relative to the TSS of the *SlDFR* promoter (gRNA:SlDFR). The left side of the leaf transiently expresses a luciferase reporter driven by the *SlDFR* promoter as a negative control of induction (no gRNA). WT represents a non-transgenic plant that transiently expresses the dCasEV2.1 components. Asterisks indicate Student’s t-test significant values (*P* < 0.05) compared with the control samples. L3, L4, L6, L7, L8, and L9 are independent YFP-dCasEV T1 lines. Bars represent average relative transcriptional activities (RTAs) ± SD, n = 3. Images generated with BioRender.com

### Engineering PVX-based sgRNA delivery for systemic gene activation

Once the CRISPRa chassis was developed, it was necessary to optimize the gRNA delivery for gene activation. Taking the previously reported work by Uranga et al. (2021a) as a starting point, a new version of the pPVX-based gRNA delivery vector was designed (PVX_VIGR vector), changing the native gRNA scaffold for the so-called gRNA2.1 scaffold, which incorporates two RNA aptamers that recognize the MS2 phage coat protein at the 3’ end of the scaffold. Also, a multiplexing structure harbouring two tandemly arrayed gRNAs under the control of the PVX CP promoter was incorporated into the recombinant pPVX viral vector (Figure 2A). Next, the efficiency of PVX_VIGR as an agroinfection-mediated delivery agent for gRNAs in systemic gene activation was tested, using the CRISPRa-ready dCasEV_L6 *N. benthamiana* line as recipient chassis. For this assay, we targeted the endogenous *NbDFR* gene, whose strong and highly specific responsiveness to dCasEV2.1 activation was earlier reported (Selma et al. 2019). Two *NbDFR*-specific gRNAs were designed for this purpose, targeting the positions -145 and -198 bp relative to the TSS of the *NbDFR* gene, and cloned into the PVX_VIGR vector, generating the PVX::gDFR construct. In a first approach, the PVX_VIGR-encoded gene activation program was agroinoculatedin basal *N. benthamiana* leaves, employing an optical density (OD) at 600 nm of 0.5 as was described previously in Uranga et al 2021a. The activation effects were recorded in the first and second symptomatic leaves, 9 and 14 days post-inoculation (dpi), respectively. The symptoms are easily noticeable due to the characteristic phenotypic alterations generated in the leaves. As shown in Figure 2B, strong transcriptional activation of *NbDFR* was observed in symptomatic leaves in plants treated with PVX::gDFR construct, but not in those treated with a PVX_VIGR vector carrying control gRNA. No significant differences in the activation rates were found between the two analysed symptomatic leaves, but a considerable increase in transcript levels was observed over time, reaching activation levels of 45- to 55-fold at 9 dpi and 250- to 370-fold at 14 dpi. The assay was repeated including a 20 dpi time point; however, no significant increase in the activation rates was observed in this time point (Supplemental Figure 1).

**Figure 2:**
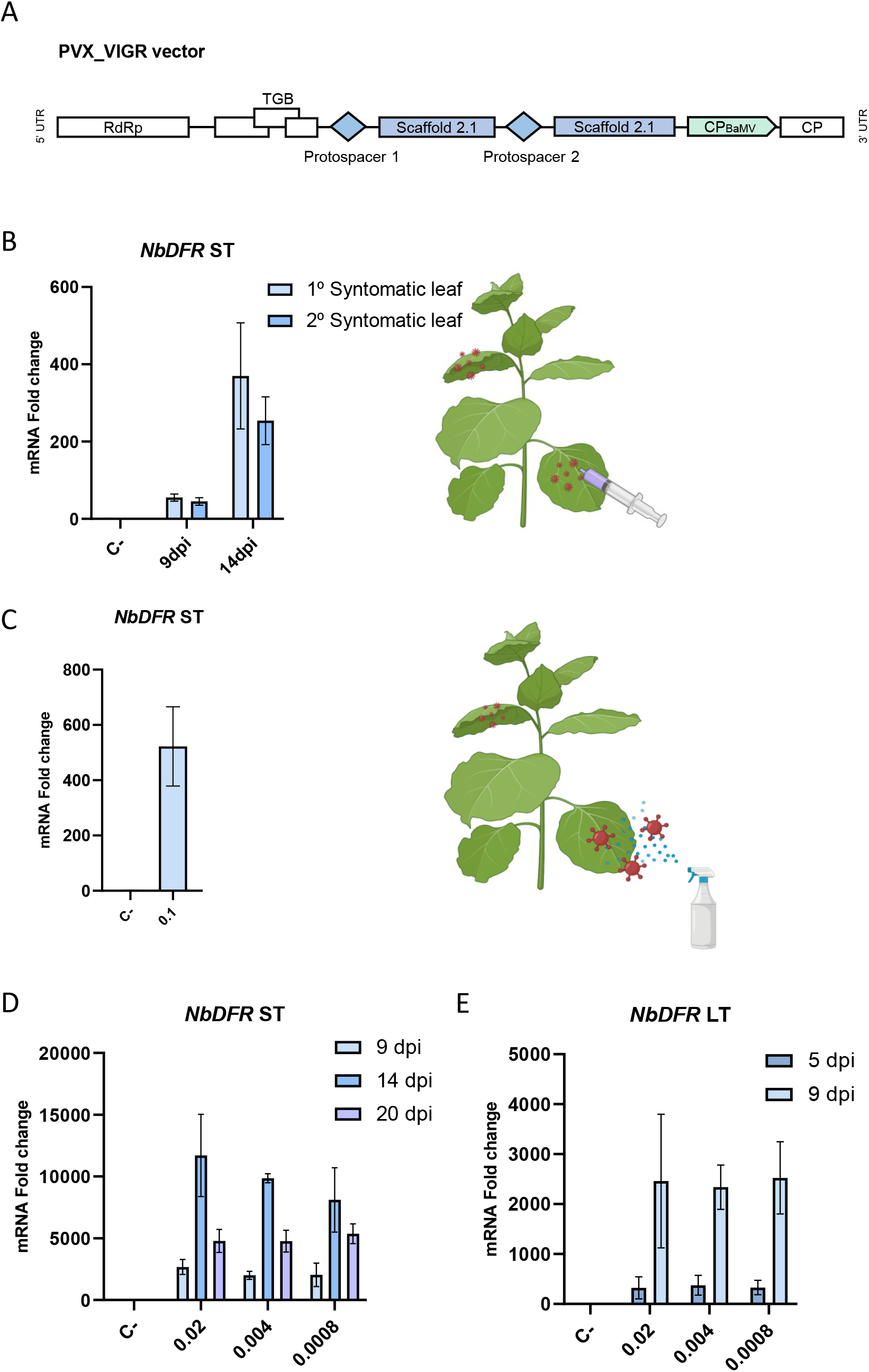
Engineering and optimization of PVX_VIGR vector for gRNA delivery. A) Schematic representation of recombinant PVX_VIGR with the gRNA2.1. The gRNA2.1 is a modification of the native gRNA scaffold with the addition of two RNA aptamers at 3’ that recognize the MS2 phage coat protein. B) *NbDFR* mRNA fold change at 9 and 14 dpi measured in the first and second systemic symptomatic leaves (ST, systemic tissues), after targeting *NbDFR* at positions -145 and -198 (relative to the TSS) with PVX-VIGA by syringe agro-inoculation. C) Same as in (B) but using spray inoculation at an optical density (OD) of 0.1. D) *NbDFR* mRNA fold changes observed when PVX-VIGA was inoculated at optical densities of 0.02, 0.004, 0.0008 through spray application. E) *NbDFR* mRNA fold change in local tissues (LT) at 5 and 9 dpi using same ODs as in (D). The C-represents a negative control where a non-specific gRNA was delivered with the PVX_VIGR. Bars represent average RTAs ± SD, n = 4. Images generated with BioRender.com.

Our initial results prompted us to implement further optimizations, following the logic that the usability of VIGR in, for example, large scale applications would depend on the simplicity of the delivery method and loads of the delivery agent required. For that reason, a spray-based agro-infection was employed as an alternative method to classical agro-infiltration with a syringe or vacuum. This method has some advantages such as a non-invasive application and the possibility of infecting many plants at the same time with simple equipment. As a first approach, an Agrobacterium optical density (OD) at 600 nm of 0.1 was employed, spraying two leaves per plant. After 14 dpi, *NbDFR* transcript levels in systemic tissues showed a successful activation of the gene, reaching up to 520-fold in the first symptomatic leaf as compared with control plants, where non-specific gRNA was delivered with the PVX_VIGR(Figure 2C). The next analysis was focused on obtaining a minimal effective OD for activation. In this case, also the youngest sprayed leaf was analysed at 5 and 9 dpi to measure the effects in local tissues of the PVX_VGR recombinant virus. Interesting, no significant differences in gene activation were found among the optical densities of 0.02, 0.004 and 0.0008, both in systemic (Figure 2D) or local infections (Figure 2E). Surprisingly, the lowest optical densities assayed in this second experiment resulted in activation rates even higher than those obtained in assays at OD= 0.1, reaching up 11,000-fold activation at 14 dpi in the first upper symptomatic leaf. As in the previous experiment, no further increases in the activation rates were observed at 20dpi. On the contrary, a slight decrease in the *NbDFR* transcript levels was observed at the later time point, perhaps due to leaf ageing. Also, successful activation of *NbDFR* was achieved in local leaves by the spraying method, with maximum activation rates of 2500 fold found at 9 dpi. These results confirm that the spray approach is an effective, easy-to-handle, non-invasive method to deliver to the plant the activation information contained in the PVX-VGR system.

### PVX_VGR spray activates three different *N. benthamiana* MYB transcription factors generating distinctive transcriptional responses

To evaluate the ability of PVX_VGR to activate endogenous genes other than *NbDFR*, we designed specific activation gRNAs programs for three endogenous MYB transcriptional factors (TFs), namely *NbODO1* (NbD050495.1), *NbMYB21* (NbD050797.1) and *NbMYB24* (NbD027779.1). The criteria for the selection of these three factors were that they (i) show high levels of expression in at least one tissue (flowers) but low levels in the leaves, and (ii) show homology with MYB factors from other species involved in the regulation of secondary metabolic profiles. With the first criterium, we wanted to ensure the inducibility of the selected genes, avoiding targeting pseudogenes or constitutively inactivated gene homeologues. With the second criterium, we aimed to show the possibility to create distinctive MYB-specific metabolite fingerprints in leaves using PVX_VGR (Figure 3A). Highly homologous genes for all three TFs were earlier described as modifiers of the phenylpropanoid composition and/or of the volatile profiles in Arabidopsis, petunia and tomato, respectively (Verdonk et al. 2005; Dal Cin et al. 2011; Huang et al. 2020). Very little is known, however, about their role in N. benthamiana.

**Figure 3:**
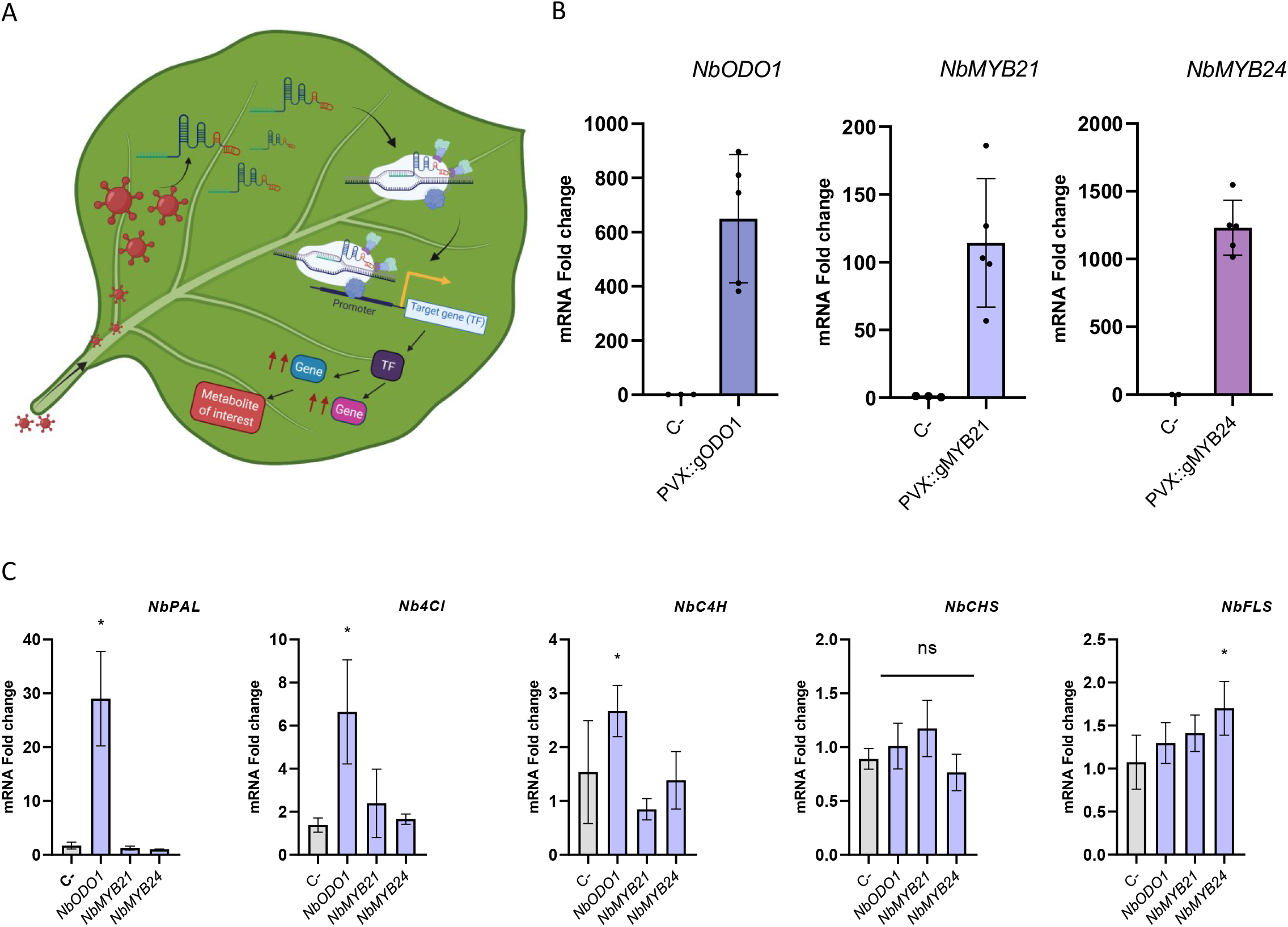
PVX_VIGR gRNA delivery for systemic activation of TFs. A) Schematic representation of the PVX_VIGR vector employed for altering the metabolite profiles. B) mRNA fold change at 14 dpi in the first symptomatic leaf (ST, systemic tissue) upon targeting the TFs *NbODO1, NbMYB21, NbMYB24* with dCasEV2.1 by spraying the PVX_VIGR vectors at an optical density (OD) of 0.008. C) mRNA fold change of *NbPAL, NbC4H, Nb4Cl, NbCHS, NbFLS* genes at 14 dpi in the first symptomatic leaf in plants where the *NbODO1, NbMYB21*, or *NbMYB24* were activated with dCasEV2.1 employing a spray inoculation of PVX_VIGR vectors at an OD of 0.008. The C-represents negative controls where a non-specific gRNA was delivered with the PVX_VIGR. Asterisks indicate Student’s *t-test* significant values (*P* < 0.05) compared with the control samples. Bars represent average RTAs ± SD, n = 5 in TFs induction analysis and n = 3 in *NbPAL, NbC4H, Nb4Cl, NbCHS, NbFLS* transcriptomic analysis. Images generated with BioRender.com.

Two gRNAs were designed for activating each of the target genes. The criteria for protospacer selection were: (i) location between -100 and -300 bp upstream of the TSS (Pan et al. 2021a), (ii) optimal on-target score, and (iii) absence of putative off-targets in the *N. benthamiana* genome. Following the same multiplexing strategy described above, the gRNAs were incorporated in the PVX_VGR vector and delivered with the spray-based method using an OD (600 nm) of 0.0008. This concentration was chosen since no differences were found after comparing the activation rates achieved with more concentrated PVX_VGR *Agrobacterium* suspensions. Next, the transcriptional activation of the targeted TFs was analysed at 14dpi in the first symptomatic leaves by RT-qPCR. The results in Figure 3B indicate that all three PVX-delivered programs resulted in significant activation of the target genes, which are otherwise repressed in leaves, reaching activation rates of >600-fold for *NbODO1*, >110 fold for *NbMYB21*, and >1200-fold for *NbMYB24* transcripts, respectively. As an indirect indication of the absolute mRNA levels obtained, the Ct values of all three TFs exceeded those observed for the F-box gene employed as a normalizing reference (between 25-26 Ct, Supplemental Table 4). Next, we analysed the changes in the transcript levels of putative downstream-regulated genes in the phenylpropanoid pathway: *NbPAL, NbCHS, Nb4CL, NbC4H*, and *NbFLS* (Battat et al. 2019; Boersma et al. 2021). As shown in Figure 3C, only the *NbODO1* activation program resulted in a strong upregulation of the *NbPAL, Nb4CL*, and *NbC4H* genes, whereas the *NbMYB24* activation program only generated a slight increase in *NbFLS*, and *NbMYB21* did not result in significant activation in any of the analysed genes.

### PVX_VGR sprayed leaves show target-specific metabolic profiles

Once the distinctive transcriptomic responses produced by each PVX_VIGR activation program were demonstrated, we decided to investigate if this had a reflection on the leaf metabolite composition, yielding three distinctive metabolite profiles. For this, we first analysed the volatile profiles of PVX_VGR-treated leaves. It shown in Figure 4A, although no drastic differences were found in the volatile profiles when the 20 most significatively different volatiles were displayed in a hierarchical cluster. In general, a significant decrease of some monoterpenes such as alpha-terpineol was observed in all the MYB-activated samples compared with PVX_VGR-treated control samples carrying gRNAs with no match in the genome. In upregulation terms, the greatest changes were found in the *NbMYB21*-sprayed plants. Here methoxycalamenene (a sesquiterpene) showed a 2,5-fold increase, and other compounds with unknown biological functions, such as 3,4,4-trimethyl-2-cyclopentene-1-one, 2,4-diisopropylphenyl acetate, and phenol 3,5-dimethyl, showed *NbMYB21*-specific overaccumulation (Figure 4B).

**Figure 4:**
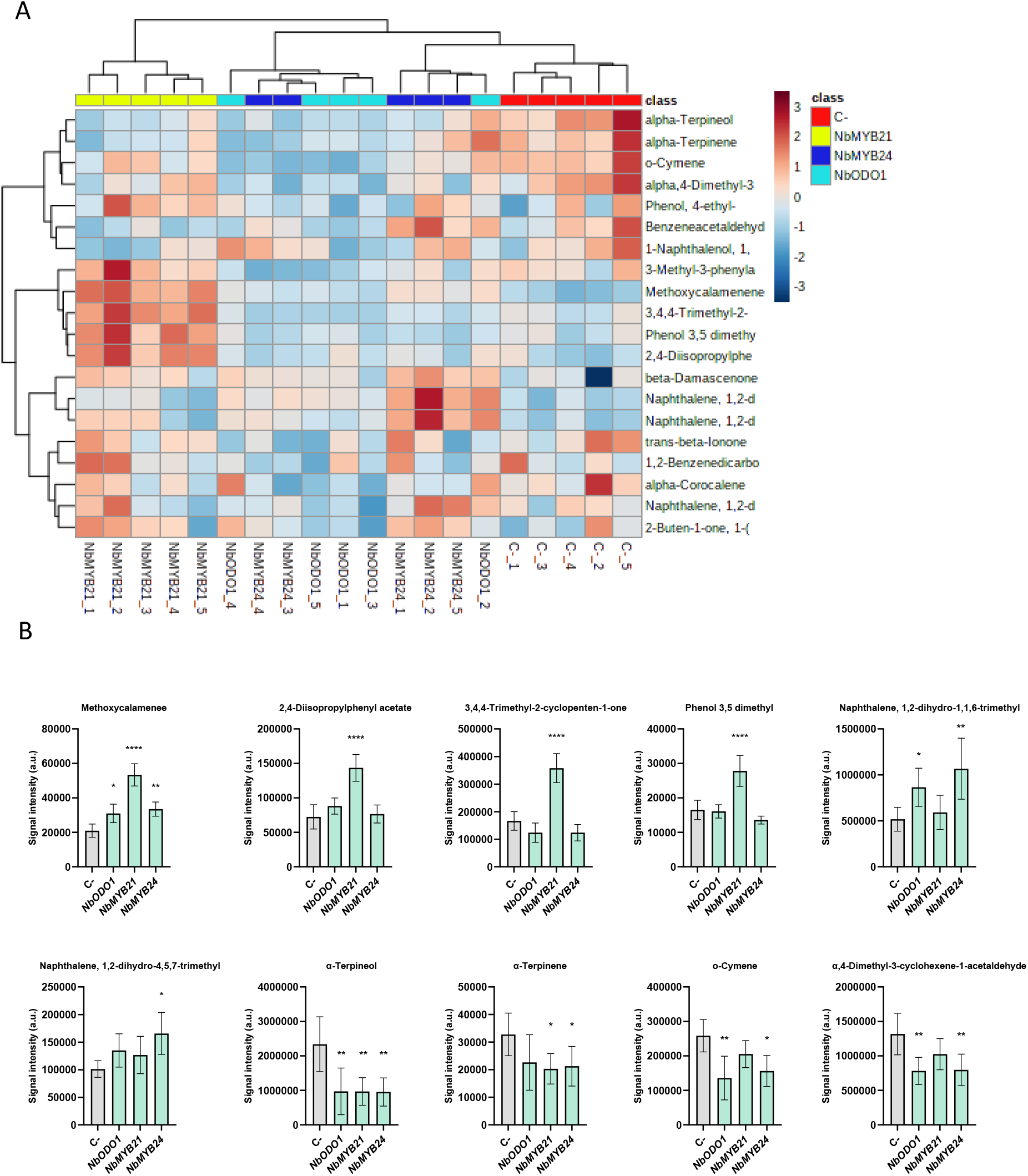
Analysis of the volatile profiles of *N. benthamiana* leaves activated in *NbODO1, NbMYB21* and *NbMYB24* expression. A) Hierarchical cluster analysis and heatmap representation of the volatile metabolite profiles obtained by GC-MS from leaves treated with PVX_VIGR vectors targeting *NbODO1, NbMYB21* and *NbMYB24*. Control samples (C-) were treated with a PVX_VIGR vector carrying a non-specific gRNA. The cluster shows the 20 most significantly different metabolites using a Student’s *t-*test analysis (*P* < 0.05). The data were obtained using Euclidean distance and Ward’s minimum variance method. Red indicates up-regulated, and blue indicates down-regulated metabolites. Five biological replicates were employed per condition. B) Signal intensity analysis of the differential compounds found in at least one of the PVX_VIGR treatments (*P*<0.05). Bars represent average fold change +/- SD (n = 5). Asterisks indicate Student’s t-test significant values (* = *P*<0.05, ** = *P*<0.01 and **** = *P*<0.0001).

The modest differences observed in the non-targeted analysis of the volatile compounds prompted us to evaluate the non-volatile metabolic profiles. For that, an untargeted LC-MS analysis was carried out with the same samples previously analysed by RT-qPCR and GC-MS. Interestingly, LC-MS revealed treatment-specific metabolic fingerprints, as the principal component analysis (PCA) in Figure 5A shows. The class-specific differences are depicted in Figure 5B where the 350 most significantly different features are hierarchically clustered. The transcription factor that generates the most distinctive non-volatile metabolic fingerprint is *NbMYB24*, followed by *NbODO1*. In contrast, *NbMYB21* activation shows a lower ability to alter the metabolite content of *N. benthamiana* leaves. A tentative identification of differential compounds, performed by the differential m/z ions and their respective retention times, was carried out as listed in Table 1 and summarized in Figure 5C. As expected, most perturbations corresponded to metabolites in the phenylpropanoid pathway. This is reflected in the increased levels of phenylalanine in *NbODO1*- (1.6 fold) and *NbMYB24*- (1.9 fold) induced samples, and in the increased levels of coumaroyl-*D*-quinic acid, the product of the *PAL, C4H* and *CL4* gene activities, in the samples sprayed with *NbODO1* (2.3fold) and *NbMYB24* (2.7 fold) activation programs. The accumulation of 5-O-caffeoyl shikimic acid and caffeic acid 3-glucoside in the *NbMYB24* (5.1and 2.2-fold respectively) and *NbODO1* samples (4.4 and 2.3-fold respectively) indicates that the activation of these two TFs leads to the upregulation of the caffeic acid branch of the phenylpropanoid pathway. On the contrary, a clear downregulation of chlorogenic acid is observed in *NbMYB21* samples (0.3-fold). Furthermore, a general reduction in N-caffeoylputrescine was observed in all three TF activated samples. Finally, in *NbODO1* samples, the metabolic flux of phenylpropanoid derivatives presents a strong shift towards lignans accumulation, as evidenced by the high accumulation of syringaresinol β-D-glucoside (> 22-fold) and sinapoylglucose (> 4-fold). Slight increases in these compounds can also be observed in *NbMYB24* and *NbMYB21* samples but to lower extent. Remarkably, flavonols over-accumulate in all TF-targeted samples, specifically kaempferol glucosides, with the highest accumulation observed in *NbODO1* samples (11-fold). Finally, other metabolic pathways also seem perturbed, although to a lower extent. A significant increase in phospholipids (Linoleoylglycerol and LysoPC), and a decrease in hydroxy steroid derivatives (esthylestrenol) were found in *NbMYB24*-induced samples.

**Table 1:**
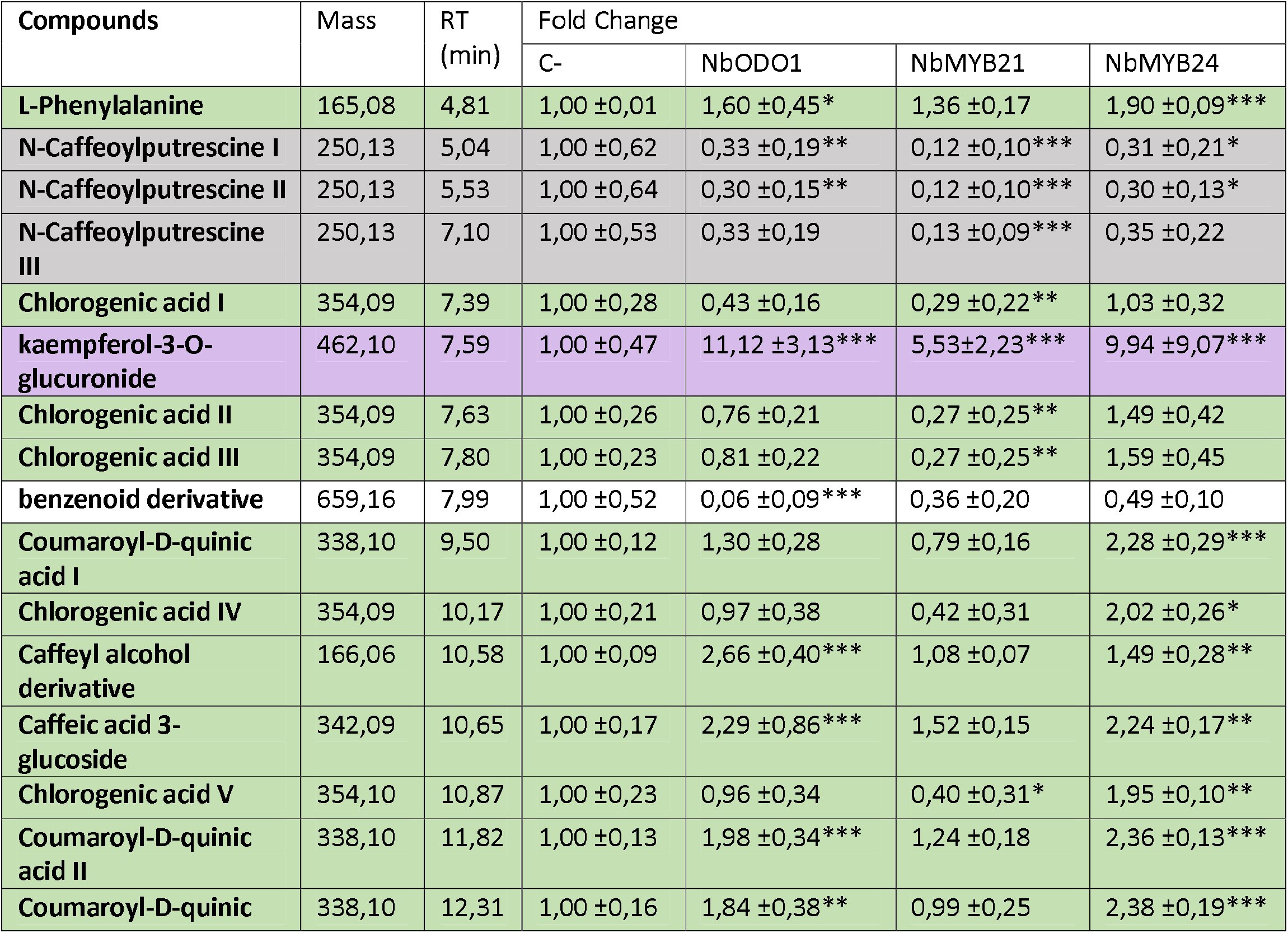
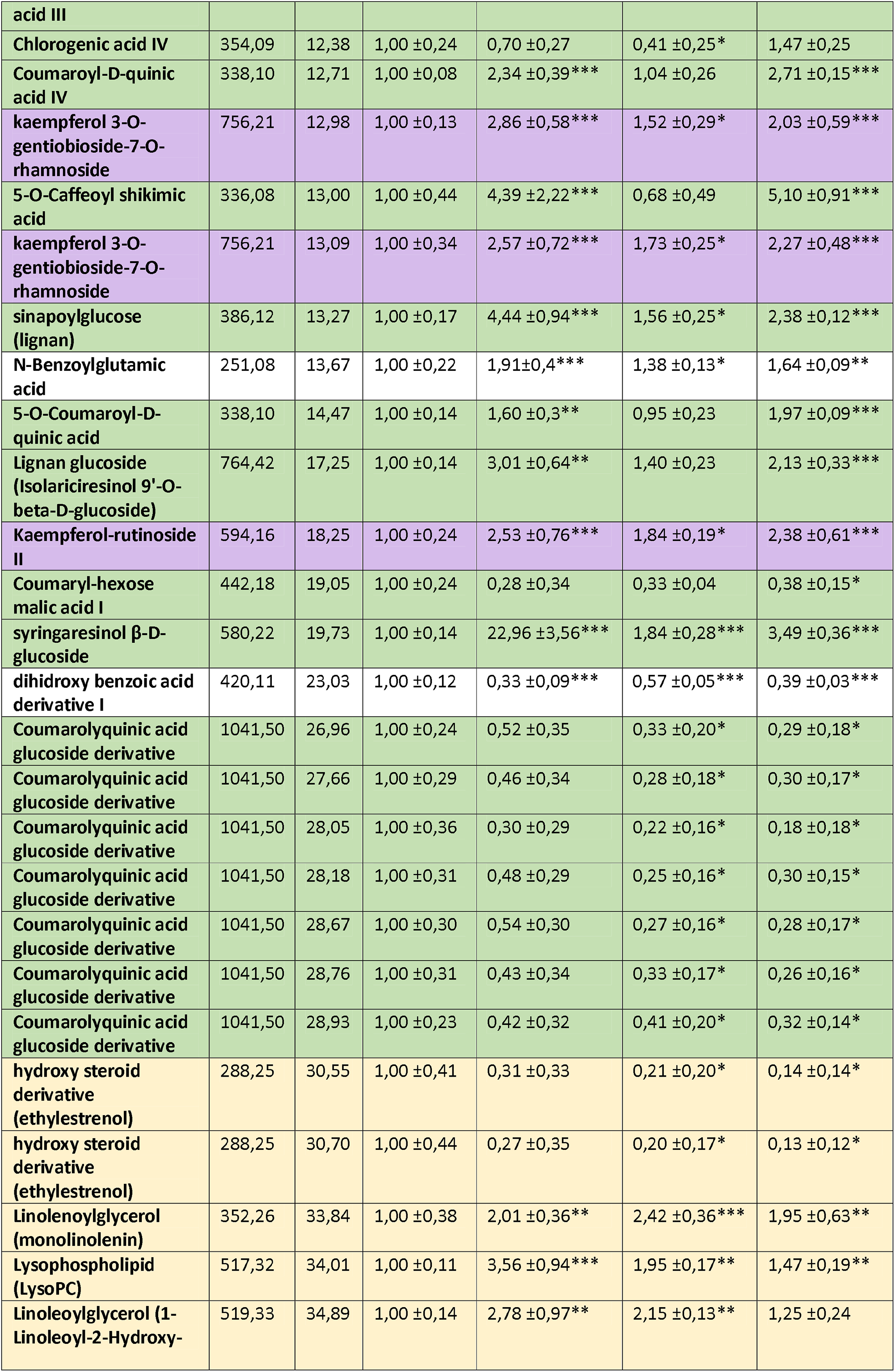
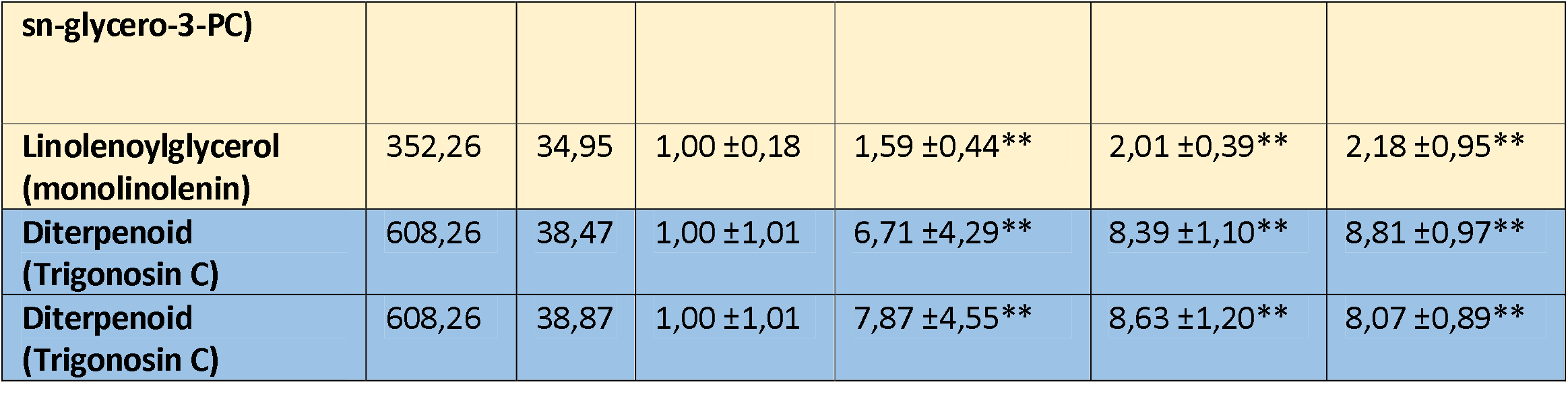
Differential tentatively identified compounds obtained from non-targeted LC-MS metabolic analysis of the leaves treated with PVX_VIGR targeting *NbODO1, NbMYB21* or *NbMYB24*. The *m/z* signal intensity values employed for quantification are normalized with the negative control (C-) values. The colouration of table cells for each compound corresponds to the colours employed in the diagram of the metabolic routes in Figure 5C. Asterisks indicate Student’s t-test significant values (* = *P*<0.05, ** = *P*<0.01 and **** = *P*<0.0001).

**Figure 5:**
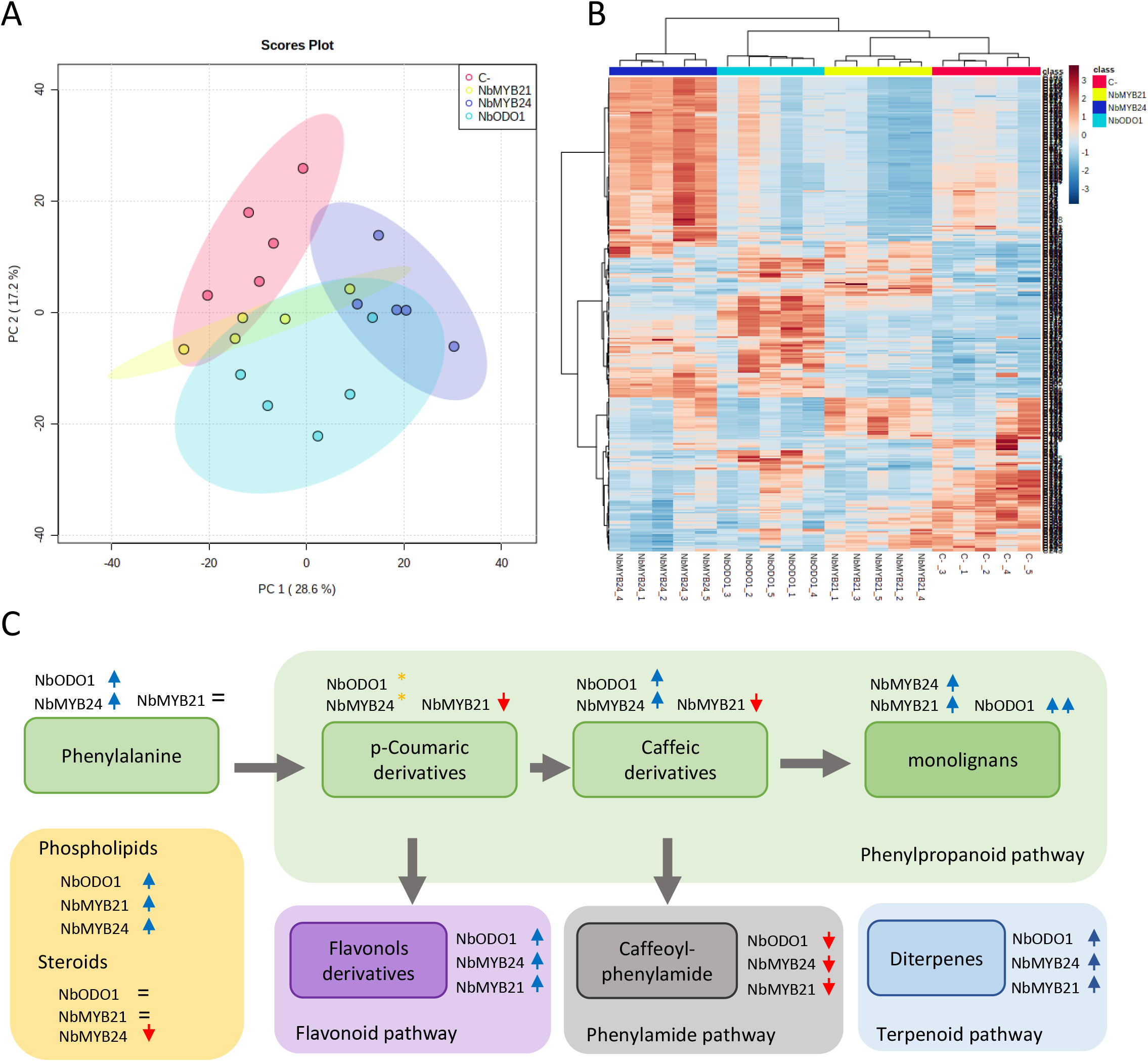
Analysis of the metabolite profiles of *N. benthamiana* plants sprayed with PVX_VIGR for targeting *NbODO1, NbMYB21* and *NbMYB24*. (A) Principal component analysis and (B) Hierarchical cluster analysis and heatmap representation resulting from the non-targeted LC-MS data obtained from leaves treated with PVX_VIGR targeting *NbODO1, NbMYB21*, or *NbMYB24*. Control samples (C-) were treated with a PVX_VIGR vector carrying a non-specific gRNA. Five biological samples were employed per condition. The *m/z* features represented in the heatmap are the 350 most significantly different using an ANOVA test (*P* < 0.05). The data were obtained using Euclidean distance and Ward’s minimum variance method. Red indicates up-regulated and blue down-regulated features. C) Schematic representation of the metabolic pathways and metabolite sub-groups that show differential abundance after inducting the *N. benthamiana* plants with a PVX-VIGR for targeting *NbODO1, NbMYB21*, or *NbMYB24* TFs compared with control samples (C-). The blue arrows represent an increment of the indicated metabolite group in the samples where *NbODO1, NbMYB21* or *NbMYB24* were up-regulated with dCasEV2.1. The red arrows represent a down-regulation of the indicated metabolite group in the samples where *NbODO1, NbMYB21*, or *NbMYB24* were up-regulated with dCasEV2.1. The orange asterisk represents that some metabolites that belong to the indicated chemical class are increased and other metabolites are down-regulated in the samples where *NbODO1, NbMYB21*, or *NbMYB24* were up-regulated with dCasEV2.1. The = symbol represents no differential metabolite changes in the sample where one of the indicated MYB TF was up-regulated with dCasEV2.1. The metabolites employed to generate this illustration were extracted from Table 1 where the differential metabolites tentatively identified are included.

## Discussion

Viral systems have been widely used in plant sciences for a variety of applications, from recombinant protein production to reverse genetics. Virus-induced gene silencing (VIGS) has been extensively used for interrogating gene function in diverse species and as a shortcut to gene knockouts bypassing stable transformation. In VIGS strategies, viral clones incorporate sequences matching endogenous genes that, consequently, induce post-transcriptional gene silencing in agro-infected plants. More recently, virus-induced gene editing (VIGE) has emerged as an alternative to transgenics for efficient gene editing. In this latter case, the gene edition machinery is usually split into two parts, a constant module comprising the Cas enzyme and a variable module comprising the gRNA. The gRNA module is encoded in the viral vector whereas the constant module, larger in size, can be either stably integrated into the plant genome (Uranga et al. 2021a) or delivered in a separate viral vector accepting larger cargo (Uranga et al. 2021b). Different viruses have been adapted for both VIGS and VIGE strategies. Whereas for VIGS, the TRV seems the ideal choice given its wide host range, mild silencing suppression, and consequently, strong gene silencing effect, in VIGE other factors need to be taken into consideration, such as the gRNA accumulation levels, the ability to self-process multiplexed gRNA arrays, and the cargo capacity, all of which also play a role in editing efficiency. Therefore, other viruses such as PVX or TMV present advantageous characteristics for VIGE (Cody et al. 2017; Ariga et al. 2020).

Similar to VIGS and VIGE, VIGR represents a new promising transient tool for plant biotechnology, with the ability not only to suppress but also to strongly induce endogenous gene expression. Thus, VIGR offers a panoply of new applications that range from gene discovery to transient genetic reprogramming of crops at a large scale, as recently proposed by Torti et al. (2021) for PVX and TMV-based transgene expression. VipariNama, a VIGR strategy based on TRV, was pioneer in demonstrating the possibilities of virus delivery in CRISPR-based transcriptional regulation in plants (Khakhar et al. 2021). Despite the elegant display of alternative VipariNama configurations, which includes both activation and repression strategies, actual gene activation/repression levels observed with VipariNama tools were relatively modest (between 2 and 5-fold), probably due to a limited intrinsic regulatory ability of the combination of TRV-encoded gRNA and Cas9-based synthetic transcription factors employed. Whereas these activation/repression levels were sufficient to induce measurable developmental and/or metabolic changes for some regulatory genes, there is certainly room for improvement if robust responses are required in large-scale applications. Our results here show that the PVX_VIGR system can induce a very strong gene activation in N. benthamiana, both local and systemic, opening the way to new and more robust applications. In this regard, a strong VIGA effect not only serves in research to elucidate gene function, but in the case of N. benthamiana, VIGA can also be envisioned as a tool for shaping plant metabolic content. Recently our group showed how CRISPR-based programable transcriptional factors can be employed to channel metabolic fluxes toward the accumulation of selected metabolites in *N. benthamiana* leaves (Selma et al. 2021). In that work, gRNA-encoded activation programs were delivered in the form of multiplex T-DNA constructs via agroinfiltration, a strategy that limits the scalability of the approach. This limitation could be circumvented with the advantages offered by VIGR. In particular, the spraying strategy described here offers a solution for large-scale delivery of gene reprogramming instructions to *N. benthamiana* biofactories, with possibilities to extend the technology to other crops given the broad host range of PVX. In principle, in VIGR strategies, transcriptional activation/repression will not be sustained in the next generations without re-inoculating the virus. However, the recent advances in CRISPR-based epigenetic modifications open new perspectives on the ability to induce epigenetic heritable traits using VIGR (Papikian et al. 2019; Ghoshal et al. 2020; Selma and Orzáez 2021).

The PVX recombinant virus presents an architecture that allows a multiplexing strategy for expressing the gRNAs without the necessity of pre-tRNAs, as was previously demonstrated by Uranga et al. 2021a. The addition of activation aptamers in the gRNA scaffold that are recognized by the MS2 protein in the recombinant virus structure has not been an obstacle to expressing active gRNAs. A possible constraint of PVX_VIGR is that cargo limitations restrict the complexity of the gene activation/repression programs that can be delivered to the plant. In metabolic reprogramming using T-DNA delivery via agroinfiltration, we previously employed up to fifteen tandemly arrayed gRNAs to successfully redirect a metabolic pathway (Selma et al. 2021). However, it is unlikely that such large gRNA tandem repeats could work as efficient programs in PVX_VIGR. Future works would need to establish these limitations and circumvent them with other strategies such as the co-delivery of non-competing viruses.

The capacity of PVX_VIGR to activate genes in *N. benthamiana* was evaluated by targeting the transcriptional factors *NbODO1, NbMYB21*, or *NbMYB24* and analysing the specific changes in the metabolite profiles in the induced plants. In this case, although the selected targets are predominantly expressed in flowers, their role in *N. benthamiana* remains uncharacterized, giving added value to the employment of the PVX_VIGR to analyse the metabolite fingerprint of each TF. The MYB factors described previously as putative orthologues of our target genes were studied in petunia, Arabidopsis, and tomato. The over-expression of these genes produced changes in the volatile and non-volatile metabolic profiles in these species. In petunia, the overexpression of PhODO1 was linked to the production of compounds involved in the scent, such as benzenoids, phenylpropanoids, fatty acids, and terpenoid derived volatiles (Verdonk et al. 2005). In addition, the petunia orthologs of MYB21 and MYB24, known as EOBI and EOBII (EMISSION OF BENZENOIDS), respectively, are also involved in the production of the components of the scent through tight crosstalk between them and PhODO1 (Spitzer-Rimon et al. 2012). In tomato, the ODO1 overexpression is also linked to the volatile production of benzenoids and phenylpropanoids, however, the biggest changes found in the tomato metabolome associated with ODO1 were the increase of non-volatile phenylpropanoid derivatives, which includes some flavonoids, especially flavonols (Dal Cin et al. 2011; Xie et al. 2016). Concurrently, in Arabidopsis, the increase of flavonols in flowers was directly correlated with the expression *of AtMYB99* (a homologue of *ODO1*), *AtMYB21*, and *AtMYB24* (Battat et al. 2019; Zhang et al. 2021a). The crosstalk among all three paralogues seems to take place similarly in petunia and Arabidopsis. However, the metabolite profiles resulting from this crosstalk are different in the two species, suggesting that in each species these genes regulate the different branches of the phenylpropanoid pathway differently. In addition, in Arabidopsis, AtMYB21 and AtMYB24 are known to interact with genes in the jasmonate pathway, leading to an increase in the production of terpenoids (Cheng et al. 2009). The upregulation of MYB homologues in *N. benthamiana* leaves using PVX_VIGR did not lead to dramatic changes in the volatile profiles, with the exemption of an increase in the levels of sesquiterpene methoxycalamenene observed in AtMYB21 samples. A probable explanation is a limited capacity of *N. benthamiana* leaves to produce volatile compounds, which may be related to a high activity of endogenous glycosyl transferases producing non-volatile glycosylated derivatives (Herpen et al. 2010; Louveau and Osbourn 2019). On the contrary, highly specific non-volatile metabolic profiles were observed associated with each TF transcriptional activation. In all three treatments, most changes observed were associated with the phenylpropanoid pathway. This correlates with previous observations in Arabidopsis and tomato, where most changes associated with ODO1, MYB21 or MYB24 overexpression were in the non-volatile fraction, specially phenylpropanoid derivatives (Xie et al. 2016; Battat et al. 2019). In particular, we found that all three treatments resulted in the overaccumulation of flavonols. On the contrary, the accumulation of other phenylpropanoids seemed dependent on the activation program applied. For example, p-coumaric acid and caffeic acid derivatives were enriched in *NbMYB24* and *NbODO1* activation programs but downregulated in *NbMYB21*-induced samples. In the case of *NbODO1*, this observation correlates with the upregulation of NbPAL and NbC4H genes observed in the RT-qPCR analysis. For *NbMYB24*, we did not observe a significant upregulation of the same genes. The observed metabolite accumulation could be explained by the activation of a set of homologous genes different from those evaluated by RT-qPCR, or by a different timing in gene activation by *NbMYB24*. Other phenylpropanoid derivatives such as monolignans, although upregulated in all three treatments, resulted in much higher accumulation in *NbODO1* samples. These differential accumulations observed in certain groups of related metabolites, such as lignans or flavonols, suggest the distinctive specificities of each MYB factor in the activation of the different genes in the pathway and demonstrate the power of PVX_VIGR in reprogramming metabolic fluxes.

In conclusion, we show that PVX-based gRNA delivery is an effective system for transient CRISPR activation. In this work, we demonstrate that plant metabolic profiles can be customized through a spray agro-inoculation of a PVX vector, expanding the current toolbox of viral systems employed for transcriptional activation. The precise and scalable spatiotemporal control of gene transcription offered by this new tool opens the door to increasingly sophisticated designs in plant synthetic biology.

## Methods

### Generation and selection of *N. benthamiana* dCasEV lines

The *N. benthamiana* YFP-dCasEV lines were generated following the transformation protocol described previously (Clemente 2006). The construct (GB2618) was transferred to the LBA4404 A. tumefaciens strain used for plant transformation. Murashige and Skoog (MS) plates supplied with kanamycin at 100 mg/L were used to select the transgenic T0 lines. In addition, YFP analysis was carried out to double-check the construct incorporation. Transgenic lines were sorted based on the T1 plants capable of generating efficient transcriptional activation. pDFR:Luciferase was employed as a reporter for testing the activation rates obtained incorporating the corresponding gRNA to target SlDFR promoter (gRNA:SlDFR). A wild type *N. benthamiana* was infiltrated in parallel with the same reporter, the gRNA:SlDFR, and dCasEV2.1 constant module construct for a fair comparison of the activation results obtained. Also, the T1 plants were evaluated by segregation analysis for selecting a single-copy T-DNA insertion.

The candidate single copy T2 of L3 and L6 lines were analysed for homozygosity and finally, YFP-dCasEV-L6.4.1 was selected as a homozygous population and kept for the following experiments. *N. benthamiana* transgenic lines were grown under 24°C/20°C light/darkness with a 16h/8h photoperiod in a growth chamber.

### gRNAs design and viral vector construction

Initially, the *N. benthamiana* endogenous gene *NbDFR* was selected for CRISPRa optimization. The protospacer was previously described in Selma et al 2019. The design of gRNAs for *NbODO1, NbMYB21*, and *NbMYB24* was performed employing the Benchling CRISPR application tool (https://www.benchling.com/). The protospacer sequences were selected within the activation window located between -100 and -300 bp upstream of the TSS for each gene (Pan et al. 2021a). Protospacer sequences are listed in Supplementary Table 1. Plasmid pPVX was previously described in Uranga et al. 2021a. The full-length PVX cDNA (GenBank accession number MT799816) is flanked by the *CaMV 35S* promoter and the *A. tumefaciens Nopaline synthase (Nos)* terminator. The PVX *CP* promoter drives the expression of the gRNA, while the viral *CP* that harbours a deletion of 29 initial codons (Dickmeis et al. 2014) is expressed under the control of a heterologous promoter derived from the *Bamboo mosaic virus* (BaMV). The double gRNA2.1 for CRISPRa was included in recombinant virus plasmids through PCR amplification with high-fidelity Phusion DNA polymerase (Thermo Scientific, Waltham, MA, USA) and Gibson DNA assembly (NEBuilder HiFi DNA assembly master mix, New England Biolabs, Ipswich, MA, USA). Primers employed for gRNA2.1 construction and adaptation for PVX recombinant plasmids are listed in Supplementary Table 2. All the plasmids were confirmed by Sanger sequencing. As negative control of infection, the PVX::gXT2 (un-specific for activation) was employed without adding the 2.1 aptamers.

### Plant inoculation

Transgenic *N. benthamiana* dCasEV2.1 plants were grown in growth chambers at 25°C under a 16/8-h day-night cycle. The inoculation was performed 4 weeks post sowing. The PVX_VIGR vectors were transferred to *A. tumefaciens* strain GV3101 by electroporation. Inoculation was carried out with overnight-grown bacterial cultures. The cultures were pelleted and resuspended on agroinfiltration solution (10 mM MES, pH 5.6, 10 mM MgCl2, and 200 μM acetosyringone) and incubated for 2 h at room temperature with agitation. The optical density of the bacterial cultures was adjusted to 0.5 at 600 nm for the syringe inoculation method and 0.1, 0.02, 0.004 and 0.0008 for the spray inoculation method. In the spray inoculation method, 0.04% of surfactant was added. The final optical density employed for the *NbODO1, NbMYB21*, and *NbMYB24* spray induction was 0.0008. Two leaves per plant were treated in both inoculation methods, using a 1ml syringe on the abaxial side of the leaf or a complete spray of both sides of the leaves. Control plants were inoculated with PVX::gXT2 following the same procedure. The samples were collected at 9 and 14 dpi from the first and second symptomatic leaf employing the syringe approach. The symptoms are easily noticeable due to the characteristic phenotypic alterations generated in the leaves. Samples from the youngest sprayed leaf were collected at 5 and 9 dpi to measure the kinetics induction of *NbDFR* in local infection. Samples from the first symptomatic leaf were collected at 9, 14 and 20 dpi to measure the kinetics of the *NbDFR* gene in systemic infection. Finally, samples from the *NbODO1, NbMYB21*, and *NbMYB24* induction assays were collected at 14 dpi from the first symptomatic leaf. All the samples were immediately frozen in liquid nitrogen and stored at −80°C.

### Luciferase/Renilla activity determination

The determination of the luciferase/renilla activity was carried out collecting one disc per leaf (d = 0.8 cm, approximately 18–19 mg) at 5 dpi. The samples were homogenized and extracted with 375µl of ‘Passive Lysis Buffer,’ followed by 10 min of centrifugation (14,000×g) at 4 °C. Then, the supernatant was collected as working plant extract. Fluc and Rluc activities were determined following the Dual-Glo® Luciferase Assay System (Promega) manufacturer’s protocol with minor modifications: 10µl of working plant extract, 40 µl of LARII and 40 µl of Stop&Glow Reagent were used. Measurements were made using a GloMax 96 Microplate Luminometer (Promega) with a 2-s delay and a 10-s measurement.

Fluc/Rluc ratios (RPUs) were determined as the mean value of three biological replicates coming from three independent agroinfiltrated leaves of the same plant. The RPUs were normalized to the Fluc/Rluc ratio obtained for a reference sample, that measures relative transcriptional activities (RTA)sof the evaluated promoter fused to the reporter.

### RNA isolation and RT-qPCR Gene Expression Analysis

Total RNA was isolated from 100mg of frozen leaf tissue employing an isolation kit (Gene Jet Plant Purification Mini Kit-ThermoScientific) according to the manufacturer’s instructions. RNA was treated with DNase-I Invitrogen Kit and its concentration was adjusted for cDNA reaction. 800ng of total RNA was used for cDNA synthesis using PrimeScript™ RT-PCR Kit (Takara) in 20 µL final volume according to the manufacturer’s recommendations. Expression levels were measured in four biological replicates for *NbDFR* induction assays and five biological replicates for *NbODO1, NbMYB21* and *NbMYB24* induction assays. Each biological replicate was measured in three technical replicates reactions in the presence of a fluorescent dye (SYBR® Premix Ex Taq) using the Applied biosystem QuantStudio 3 equipment. The specific primers for detection of the target genes are listed in Supplementary Table 3. F-BOX protein gene was used as an internal reference gene (Liu et al. 2012). Basal expression levels were calculated from the control samples inoculated with PVX::gXT2 recombinant plasmid. Calculations of each sample were carried out according to the comparative ^ΔΔ^ CT method (Livak et al. 2001).

### Volatile organic compound analysis and statistics

Frozen and ground symptomatic leaf samples (70 mg) were incorporated in a 10-mL headspace screw-cap vial. The samples were prepared by adding 1 mL of 5 M CaCl2, 150 μL of 500 mM EDTA (pH = 7.5) and 3 μL 10 ppm lavadulol as an internal reference and sonicated for 5 min. Volatile compounds were captured by solid-phase microextraction (HS-SPME) with a 65 μm polydimethylsiloxane/divinylbenzene (PDMS/DVB) SPME fibre (Supelco, Bellefonte, PA, USA). Volatile extraction was performed automatically by a CombiPAL autosampler (CTC Analytics). Vials were first incubated at 80°C for 3 min with 500 rpm agitation. Then, the fibre was exposed to the vial through the headscape for 20 min under the same conditions of temperature and agitation. Desorption was performed at 250°C for 1 min (splitless mode) in the injection port of a 6890 N gas chromatograph (Agilent Technologies). After desorption, the fibre was cleaned in an SPME fibre conditioning station (CTC Analytics) at 250°C for 5 min under a helium flow. Chromatography was performed on a DB5ms (60 m, 0.25 mm, 1 μm) capillary column (J&W) with helium as the carrier gas at a constant flow of 1.2 mL min^-1^. The oven conditions were 40°C for 2 min, and a 5°C min^-1^ ramp was programmed until reaching 300°C with a final hold at 300°C for 5 min. Two blank controls with 1 mL of 5 M CaCl2 and 150 μL of 500 mM EDTA were included in the experiment.

Identification of the volatile compounds was performed by employing a customized library of *N. benthamiana* based on the NIST database (See supplementary data). A non-targeted analysis for the differential emission of volatile compounds in the *NbODO1, NbMYB21*, and *NbMYB24* samples was performed. Peak intensities were calculated by employing the Agilent Mass Hunter Workstation Software for Quantitative Analysis. Compounds with a peak intensity sample:blank ratio < 3 were removed for each condition. Peak areas were normalized using the total ion count (TIC).

The resulting compounds were evaluated through a Student’s t-test student (*P*<0.05) analysis to identify differentially accumulated compounds between the control samples and the MYB-targeted samples. The spectrum profile of the differential compounds is included in Supplementary Table 5.

### Liquid chromatography and untargeted analysis

The material used for the GC-MS analysis was further analysed by LC-MS analysis. First symptomatic leaf samples of five different plants infected with each PVX_VIGR vector were collected at 14 dpi. Frozen ground tissue (50 mg) was extracted in 75% acetonitrile in water (500 µL) with 1 ppm genistein as the internal standard. The homogenate was vortexed, sonicated for 10 min, and centrifuged for 5 min at 14,000 rpm. Supernatants were filtered with a 0.2 µM filter. The analysis comprised five biological replicates per genotype.

The LC-MS analysis was performed with an Orbitrap Exploris 120 mass spectrometer coupled with a Vanquish UHPLC System (Thermo Fisher Scientific, Waltham, MA, USA). Compounds were separated by reverse-phase ultraperformance liquid chromatography using an Acquity PREMIER BEH C18 UPLC column (1.7 µM particle size, dimensions 2.1 × 150 mm) (Waters Corp., Milford, MA, USA).

Mobile phases consisted of 0.1% formic acid in water (phase A) and 0.1% formic acid in acetonitrile (phase B). The solvent gradient program was conditioned as follows: 0.5% solvent B over the first 2 min, 0.5–30% solvent B over 25 min, 30–100% solvent B over 13 min, 2 min at 100% B, return to the initial 0.5% solvent B over 1 min, and conditioning at 0.5% B for 2 min. The flow rate was 0.4 ml/min, and the injection volume was 1 µL. The column temperature was set at 40°C.

Ionisation was performed with heated electrospray ionization (H-ESI) in positive and negative modes. Samples were acquired in full scan mode (resolution set at 120000 measured at FWHM) and mixes were acquired in both full scan and data-dependent acquisition (DDA) to help with compound identification. For DDA, the resolution was set at 30000 and the intensity threshold at 2e5. The mass range was set from 150 to 1500.

Data analysis and statistics were performed with Compound Discoverer 3.3 software (Thermo Scientific, Waltham, MA, USA) and the MetaboAnalyst5.0 Software (https://www.metaboanalyst.ca/). Logarithmic transformation and autoscaling scaling were employed as normalization to perform the principal component analysis and the hierarchical clustering and to produce a heatmap. Euclidean distance and Ward clustering algorithm were applied as parameters for making hierarchical clustering, and ANOVA test was the statistical method used for generating the list of 350 molecular features with significantly different accumulation. The molecular features determined as significantly different and tentatively identified through Kegg databases associated with the Compound Discoverer 3.3 software were manually added to Table 1. The quantification of the metabolites was performed employing the parental ion and normalized with the negative control samples (C-), where an unspecific non-specific gRNA was delivered with the PVX_VIGR.

## Supplementary data

The datasets and chromatograms generated in this work are located: https://doi.org/10.5281/zenodo.6475547

## Author contributions

D.O. and S.S. designed the experiments. S.S, G.S, and U.M conducted the experiments. G.S, A-R.A, and V-V.M. contributes to data analysis. D.O. and S.S drafted the manuscript. G.S, D.M, U.M, F.PD and D.JA contribute to the manuscript writing and editing. All the authors discussed and revised the manuscript.

## Acknowledgements

This work has been funded by Grant PID2019-108203RB-10 Plan Nacional I+D, Spanish Ministry of Economy and Competitiveness, and Spanish Ministry of Science and Innovation. Sara Selma is a recipient of FPI fellowship associated with this Grant (BES-2017-080098).

## References

Ali Z, Abul-faraj A, Li L, Ghosh N, Piatek M, Mahjoub A, Aouida M, Piatek A, Baltes NJ, Voytas DF, Dinesh-Kumar S, Mahfouz MM (2015) Efficient Virus-Mediated Genome Editing in Plants Using the CRISPR/Cas9 System. Mol Plant 8(8):1288–1291. https://doi.org/10.1016/j.molp.2015.02.011

Ali Z, Eid A, Ali S, Mahfouz MM (2018) Pea early-browning virus-mediated genome editing via the CRISPR/Cas9 system in Nicotiana benthamiana and Arabidopsis. Virus Research 244:333– 337. https://doi.org/10.1016/j.virusres.2017.10.009

Ariga H, Toki S, Ishibashi K (2020) Potato Virus X Vector-Mediated DNA-Free Genome Editing in Plants. Plant and Cell Physiology 61(11):1946–1953. https://doi.org/10.1093/pcp/pcaa123

Battat M, Eitan A, Rogachev I, Hanhineva K, Fernie A, Tohge T, Beekwilder J, Aharoni A (2019) A MYB Triad Controls Primary and Phenylpropanoid Metabolites for Pollen Coat Patterning. Plant Physiol 180(1):87–108. https://doi.org/10.1104/pp.19.00009

Boersma MR, Patrick RM, Jillings SL, Shaipulah NFM, Sun P, Haring MA, Dudareva N, Li Y, Schuurink RC (2021) ODORANT1 targets multiple metabolic networks in petunia flowers. The Plant Journal n/a(n/a). https://doi.org/10.1111/tpj.15618

Charfeddine M, Samet M, Charfeddine S, Bouaziz D, Gargouri Bouzid R (2019) Ectopic Expression of StERF94 Transcription Factor in Potato Plants Improved Resistance to Fusarium solani Infection. Plant Molecular Biology Reporter 37(5):450–463. https://doi.org/10.1007/s11105-019-01171-4

Chavez A, Scheiman J, Vora S, Pruitt BW, Tuttle M, P R Iyer E, Lin S, Kiani S, Guzman CD, Wiegand DJ, Ter-Ovanesyan D, Braff JL, Davidsohn N, Housden BE, Perrimon N, Weiss R, Aach J, Collins JJ, Church GM (2015) Highly efficient Cas9-mediated transcriptional programming. Nat Methods 12(4):326–328. https://doi.org/10.1038/nmeth.3312

Cheng H, Song S, Xiao L, Soo HM, Cheng Z, Xie D, Peng J (2009) Gibberellin Acts through Jasmonate to Control the Expression of MYB21, MYB24, and MYB57 to Promote Stamen Filament Growth in Arabidopsis. PLOS Genetics 5(3):e1000440. https://doi.org/10.1371/journal.pgen.1000440

Clemente T (2006) Nicotiana (Nicotiana tabaccum, Nicotiana benthamiana). In: Wang K (ed) Agrobacterium protocols. Springer, pp 143–154

Cody WB, Scholthof HB, Mirkov TE (2017) Multiplexed Gene Editing and Protein Overexpression Using a Tobacco mosaic virus Viral Vector. Plant Physiology 175(1):23–35. https://doi.org/10.1104/pp.17.00411

Dal Cin V, Tieman DM, Tohge T, McQuinn R, de Vos RCH, Osorio S, Schmelz EA, Taylor MG, Smits-Kroon MT, Schuurink RC, Haring MA, Giovannoni J, Fernie AR, Klee HJ (2011) Identification of genes in the phenylalanine metabolic pathway by ectopic expression of a MYB transcription factor in tomato fruit. Plant Cell 23(7):2738–2753. https://doi.org/10.1105/tpc.111.086975

Danilo B, Perrot L, Mara K, Botton E, Nogué F, Mazier M (2019) Efficient and transgene-free gene targeting using Agrobacterium-mediated delivery of the CRISPR/Cas9 system in tomato. Plant Cell Reports 38(4):459–462. https://doi.org/10.1007/s00299-019-02373-6

Dickmeis C, Fischer R, Commandeur U (2014) Potato virus X-based expression vectors are stabilized for long-term production of proteins and larger inserts. Biotechnology Journal 9(11):1369–1379. https://doi.org/10.1002/biot.201400347

Ellison EE, Nagalakshmi U, Gamo ME, Huang P, Dinesh-Kumar S, Voytas DF (2020) Multiplexed heritable gene editing using RNA viruses and mobile single guide RNAs. Nat Plants 6(6):620– 624. https://doi.org/10.1038/s41477-020-0670-y

Gentzel IN, Ohlson EW, Redinbaugh MG, Wang G-L (2022) VIGE: virus-induced genome editing for improving abiotic and biotic stress traits in plants. Stress Biology 2(1):2. https://doi.org/10.1007/s44154-021-00026-x

Ghoshal B, Vong B, Picard CL, Feng S, Tam JM, Jacobsen SE (2020) A viral guide RNA delivery system for CRISPR-based transcriptional activation and heritable targeted DNA demethylation in Arabidopsis thaliana. PLoS Genet 16(12):e1008983–e1008983. https://doi.org/10.1371/journal.pgen.1008983

Gleba YY, Tusé D, Giritch A (2013) Plant Viral Vectors for Delivery by Agrobacterium. In: Palmer K, Gleba Y (eds) Plant Viral Vectors. Springer Berlin Heidelberg, Berlin, Heidelberg, pp 155–192

Hamada H, Linghu Q, Nagira Y, Miki R, Taoka N, Imai R (2017) An in planta biolistic method for stable wheat transformation. Scientific Reports 7(1):11443. https://doi.org/10.1038/s41598-017-11936-0

Hu J, Li S, Li Z, Li H, Song W, Zhao H, Lai J, Xia L, Li D, Zhang Y (2019) A barley stripe mosaic virus-based guide RNA delivery system for targeted mutagenesis in wheat and maize. Molecular Plant Pathology 20(10):1463–1474. https://doi.org/10.1111/mpp.12849

Huang H, Gong Y, Liu B, Wu D, Zhang M, Xie D, Song S (2020) The DELLA proteins interact with MYB21 and MYB24 to regulate filament elongation in Arabidopsis. BMC Plant Biology 20(1):64. https://doi.org/10.1186/s12870-020-2274-0

Jiang W, Li H, Liu X, Zhang J, Zhang W, Li T, Liu L, Yu X (2020) Precise and efficient silencing of mutant Kras(G12D) by CRISPR-CasRx controls pancreatic cancer progression. Theranostics 10(25):11507–11519. https://doi.org/10.7150/thno.46642

Khakhar A, Wang C, Swanson R, Stokke S, Rizvi F, Sarup S, Hobbs J, Voytas DF (2021) VipariNama: RNA viral vectors to rapidly elucidate the relationship between gene expression and phenotype. Plant Physiology 186(4):2222–2238. https://doi.org/10.1093/plphys/kiab197

Lee JE, Neumann M, Duro DI, Schmid M (2019) CRISPR-based tools for targeted transcriptional and epigenetic regulation in plants. PLOS ONE 14(9):e0222778. https://doi.org/10.1371/journal.pone.0222778

Lessard PA, Kulaveerasingam H, York GM, Strong A, Sinskey AJ (2002) Manipulating Gene Expression for the Metabolic Engineering of Plants. Metabolic Engineering 4(1):67–79. https://doi.org/10.1006/mben.2001.0210

Li T, Hu J, Sun Y, Li B, Zhang D, Li W, Liu J, Li D, Gao C, Zhang Y, Wang Y (2021) Highly efficient heritable genome editing in wheat using an RNA virus and bypassing tissue culture. Molecular Plant 14(11):1787–1798. https://doi.org/10.1016/j.molp.2021.07.010

Liang Z, Chen K, Gao C (2019) Biolistic Delivery of CRISPR/Cas9 with Ribonucleoprotein Complex in Wheat. In: Qi Y (ed) Plant Genome Editing with CRISPR Systems: Methods and Protocols. Springer New York, New York, NY, pp 327–335

Lico C, Benvenuto E, Baschieri S (2015) The Two-Faced Potato Virus X: From Plant Pathogen to Smart Nanoparticle. Frontiers in Plant Science 6

Liu D, Shi L, Han C, Yu J, Li D, Zhang Y (2012) Validation of Reference Genes for Gene Expression Studies in Virus-Infected Nicotiana benthamiana Using Quantitative Real-Time PCR. PLOS ONE 7(9):e46451. https://doi.org/10.1371/journal.pone.0046451

Livak KJ, Schmittgen TD (2001) Analysis of Relative Gene Expression Data Using Real-Time Quantitative PCR and the 2−ΔΔCT Method. Methods 25(4):402–408. https://doi.org/10.1006/meth.2001.1262

Louveau T, Osbourn A (2019) The Sweet Side of Plant-Specialized Metabolism. Cold Spring Harb Perspect Biol 11(12):a034744. https://doi.org/10.1101/cshperspect.a034744

Lowder LG, Zhou J, Zhang Y, Malzahn A, Zhong Z, Hsieh T-F, Voytas DF, Zhang Y, Qi Y (2018) Robust Transcriptional Activation in Plants Using Multiplexed CRISPR-Act2.0 and mTALE-Act Systems. Molecular Plant 11(2):245–256. https://doi.org/10.1016/j.molp.2017.11.010

Lu R, Martin-Hernandez AM, Peart JR, Malcuit I, Baulcombe DC (2003) Virus-induced gene silencing in plants. Methods 30(4):296–303. https://doi.org/10.1016/S1046-2023(03)00037-9

Maeda AE, Nakamichi N (2022) Plant clock modifications for adapting flowering time to local environments. Plant Physiology :kiac107. https://doi.org/10.1093/plphys/kiac107

Mao Y, Botella JR, Liu Y, Zhu J-K (2019) Gene editing in plants: progress and challenges. National Science Review 6(3):421–437. https://doi.org/10.1093/nsr/nwz005

Marillonnet S, Thoeringer C, Kandzia R, Klimyuk V, Gleba Y (2005) Systemic Agrobacterium tumefaciens–mediated transfection of viral replicons for efficient transient expression in plants. Nat Biotechnol 23(6):718–723. https://doi.org/10.1038/nbt1094

Mathur J, Koncz C (1998) PEG-mediated protoplast transformation with naked DNA. Methods Mol Biol 82:267–276. https://doi.org/10.1385/0-89603-391-0:267

Mei Y, Beernink BM, Ellison EE, Konečná E, Neelakandan AK, Voytas DF, Whitham SA (2019) Protein expression and gene editing in monocots using foxtail mosaic virus vectors. Plant Direct 3(11):e00181–e00181. https://doi.org/10.1002/pld3.181

Molina-Hidalgo FJ, Vazquez-Vilar M, D’Andrea L, Demurtas OC, Fraser P, Giuliano G, Bock R, Orzáez D, Goossens A (2020) Engineering Metabolism in Nicotiana Species: A Promising Future. Trends Biotechnol. https://doi.org/10.1016/j.tibtech.2020.11.012

Moradpour M, Abdulah SNA (2020) CRISPR/dCas9 platforms in plants: strategies and applications beyond genome editing. Plant Biotechnology Journal 18(1):32–44. https://doi.org/10.1111/pbi.13232

Pan C, Sretenovic S, Qi Y (2021a) CRISPR/dCas-mediated transcriptional and epigenetic regulation in plants. Curr Opin Plant Biol 60:101980. https://doi.org/10.1016/j.pbi.2020.101980

Pan C, Wu X, Markel K, Malzahn AA, Kundagrami N, Sretenovic S, Zhang Y, Cheng Y, Shih PM, Qi Y (2021b) CRISPR–Act3.0 for highly efficient multiplexed gene activation in plants. Nat Plants 7(7):942–953. https://doi.org/10.1038/s41477-021-00953-7

Papikian A, Liu W, Gallego-Bartolomé J, Jacobsen SE (2019) Site-specific manipulation of Arabidopsis loci using CRISPR-Cas9 SunTag systems. Nature communications 10(1):729. https://doi.org/10.1038/s41467-019-08736-7

Platt RJ, Chen S, Zhou Y, Yim MJ, Swiech L, Kempton HR, Dahlman JE, Parnas O, Eisenhaure TM, Jovanovic M, Graham DB, Jhunjhunwala S, Heidenreich M, Xavier RJ, Langer R, Anderson DG, Hacohen N, Regev A, Feng G, Sharp PA, Zhang F (2014) CRISPR-Cas9 Knockin Mice for Genome Editing and Cancer Modeling. Cell 159(2):440–455. https://doi.org/10.1016/j.cell.2014.09.014

Selma S, Bernabé-Orts JM, Vazquez-Vilar M, Diego-Martin B, Ajenjo M, Garcia-Carpintero V, Granell A, Orzaez D (2019) Strong gene activation in plants with genome-wide specificity using a new orthogonal CRISPR/Cas9-based programmable transcriptional activator. Plant Biotechnol J 17(9):1703–1705. https://doi.org/10.1111/pbi.13138

Selma S, Orzáez D (2021) Perspectives for epigenetic editing in crops. Transgenic Research 30(4):381–400. https://doi.org/10.1007/s11248-021-00252-z

Selma S, Sanmartín N, Espinosa-Ruiz A, Gianoglio S, Lopez-Gresa M, Vázquez-Vilar M, Flors V, Granell A, Orzaez D (2021) Custom-made design of metabolite composition in N. benthamiana leaves using CRISPR activators. Synthetic Biology

Senís E, Fatouros C, Große S, Wiedtke E, Niopek D, Mueller A-K, Börner K, Grimm D (2014) CRISPR/Cas9-mediated genome engineering: An adeno-associated viral (AAV) vector toolbox. Biotechnology Journal 9(11):1402–1412. https://doi.org/10.1002/biot.201400046

Spitzer-Rimon B, Farhi M, Albo B, Cna’ani A, Ben Zvi MM, Masci T, Edelbaum O, Yu Y, Shklarman E, Ovadis M, Vainstein A (2012) The R2R3-MYB-like regulatory factor EOBI, acting downstream of EOBII, regulates scent production by activating ODO1 and structural scent-related genes in petunia. Plant Cell 24(12):5089–5105. https://doi.org/10.1105/tpc.112.105247

Tiwari SB, Belachew A, Ma SF, Young M, Ade J, Shen Y, Marion CM, Holtan HE, Bailey A, Stone JK, Edwards L, Wallace AD, Canales RD, Adam L, Ratcliffe OJ, Repetti PP (2012) The EDLL motif: a potent plant transcriptional activation domain from AP2/ERF transcription factors: Strong plant transcriptional activation domain. The Plant Journal 70(5):855–865. https://doi.org/10.1111/j.1365-313X.2012.04935.x

Torti S, Schlesier R, Thümmler A, Bartels D, Römer P, Koch B, Werner S, Panwar V, Kanyuka K, Wirén N von, Jones JDG, Hause G, Giritch A, Gleba Y (2021) Transient reprogramming of crop plants for agronomic performance. Nat Plants 7(2):159–171. https://doi.org/10.1038/s41477-021-00851-y

Tzfira T, Citovsky V (2006) Agrobacterium-mediated genetic transformation of plants: biology and biotechnology. Current Opinion in Biotechnology 17(2):147–154. https://doi.org/10.1016/j.copbio.2006.01.009

Uranga M, Aragonés V, Selma S, Vázquez-Vilar M, Orzáez D, Daròs J-A (2021a) Efficient Cas9 multiplex editing using unspaced sgRNA arrays engineering in a Potato virus X vector. Plant J 106(2):555–565. https://doi.org/10.1111/tpj.15164

Uranga M, Vazquez-Vilar M, Orzáez D, Daròs J-A (2021b) CRISPR-Cas12a Genome Editing at the Whole-Plant Level Using Two Compatible RNA Virus Vectors. The CRISPR Journal 4(5):761–769. https://doi.org/10.1089/crispr.2021.0049

van Herpen TWJM, Cankar K, Nogueira M, Bosch D, Bouwmeester HJ, Beekwilder J (2010) Nicotiana benthamiana as a Production Platform for Artemisinin Precursors. PLOS ONE 5(12):e14222. https://doi.org/10.1371/journal.pone.0014222

Verdonk JC, Haring MA, van Tunen AJ, Schuurink RC (2005) ODORANT1 regulates fragrance biosynthesis in petunia flowers. Plant Cell 17(5):1612–1624. https://doi.org/10.1105/tpc.104.028837

Wu S, Zhu H, Liu J, Yang Q, Shao X, Bi F, Hu C, Huo H, Chen K, Yi G (2020) Establishment of a PEG-mediated protoplast transformation system based on DNA and CRISPR/Cas9 ribonucleoprotein complexes for banana. BMC Plant Biology 20(1):425. https://doi.org/10.1186/s12870-020-02609-8

Wydro M, Kozubek E, Lehmann P (2006) Optimization of transient Agrobacterium-mediated gene expression system in leaves of Nicotiana benthamiana. Acta Biochim Pol 53(2):289–298

Xie Q, Liu Z, Meir S, Rogachev I, Aharoni A, Klee HJ, Galili G (2016) Altered metabolite accumulation in tomato fruits by coexpressing a feedback-insensitive AroG and the PhODO1 MYB-type transcription factor. Plant Biotechnol J 14(12):2300–2309. https://doi.org/10.1111/pbi.12583

Xu CL, Ruan MZC, Mahajan VB, Tsang SH (2019) Viral Delivery Systems for CRISPR. Viruses 11(1). https://doi.org/10.3390/v11010028

Yin K, Han T, Liu G, Chen T, Wang Y, Yu AYL, Liu Y (2015) A geminivirus-based guide RNA delivery system for CRISPR/Cas9 mediated plant genome editing. Sci Rep 5:14926–14926. https://doi.org/10.1038/srep14926

Zhang X, He Y, Li L, Liu H, Hong G (2021a) Involvement of the R2R3-MYB transcription factor MYB21 and its homologs in regulating flavonol accumulation in Arabidopsis stamen. J Exp Bot 72(12):4319–4332. https://doi.org/10.1093/jxb/erab156

Zhang Y, Iaffaldano B, Qi Y (2021b) CRISPR ribonucleoprotein-mediated genetic engineering in plants. Plant Communications 2(2):100168. https://doi.org/10.1016/j.xplc.2021.100168

Zhang Y, Ren Q, Tang X, Liu S, Malzahn AA, Zhou J, Wang J, Yin D, Pan C, Yuan M, Huang L, Yang H, Zhao Y, Fang Q, Zheng X, Tian L, Cheng Y, Le Y, McCoy B, Franklin L, Selengut JD, Mount SM, Que Q, Zhang Y, Qi Y (2021c) Expanding the scope of plant genome engineering with Cas12a orthologs and highly multiplexable editing systems. Nature Communications 12(1):1944. https://doi.org/10.1038/s41467-021-22330-w

