## Supplementary Figure 1 for "Potato virus X -delivered CRISPR activation programs lead to strong endogenous gene induction and transient metabolic reprogramming in *Nicotiana benthamiana*"

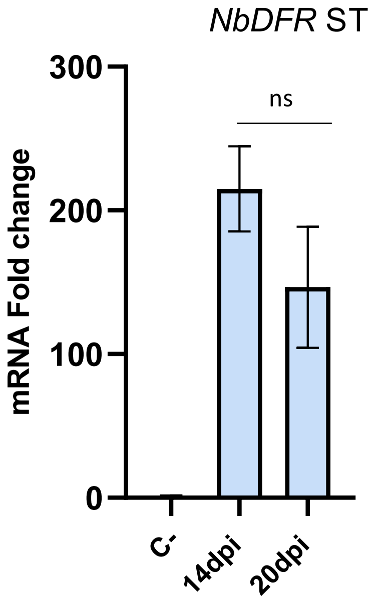


**Supplementary Figure 1: Evaluation of PVX-based sgRNA delivery for CRISPRa.** *NbDFR* mRNA fold change at 14 and 20 dpi obtained from the first symptomatic leaf (SI, systemic tissue), by targeting the endogenous gene *NbDFR* with dCasEV2.1 at positions -145 and -198 bp relative to the TSS through the PVX-based sgRNA delivery by syringe inoculation. The C- represents a negative control where an unspecific gRNA was delivered with the PVX-VIGR. Bars represent average RTAs ± SD, n = 3.
