## Supplementary tables for "Potato virus X -delivered CRISPR activation programs lead to strong endogenous gene induction and transient metabolic reprogramming in *Nicotiana benthamiana*"

| Gene | Protospacer sequence |
|---|---|
| *NbDFR* | gRNA1: ATGACTGACTGGTTGGTGAG<br>gRNA2: TATCCGTATGCCTTACCTTT |
| *NbODO1* | gRNA1: TTGGGTGGAGCTGATAACAC<br>gRNA2: TGACTAAGAAAAAAGGAAAG |
| *NbMYB21* | gRNA1: ATTTTATAGTAAAGCAAGTT<br>gRNA2: CTATTGATGTTATCATGGCA |
| *NbMYB24* | gRNA1: AATTCAGTCTTTAATCAGCC<br>gRNA2: ATTAGCAGTATATCGACTTA |

**Supplementary Table 1:** protospacer sequences designed for targeting the *N. Benthamiana* genes with dCasEV2.1

| **Construct** | **Primer sequence (5'->3')** |
|---|---|
| PVX::gDFR | Fw:gaggtcagcaccagctagcaATGACTGACTGGTTGGTGAGGTTTTAGAGCTAGAAATAGC<br>Rv: catacggataGGGAAGACTCCCCAGTGAC<br><br>Fw: gagtcttcccTATCCGTATGCCTTACCTTTGTTTTAG<br>Rv: gggaaacttaacaaaccctaGGGAAGACTCCCCAGTGAC |
| PVX::gODO1 | Fw: gaggtcagcaccagctagcaAGCATCAACAGGTCTTAAGCGTTTTAG<br>Rv: tagtgtaaatGGGAAGACTCCCCAGTGA<br><br>Fw: gagtcttcccATTTACACTAACAATAGTAGGTTTTAGA<br>Rv: gggaaacttaacaaaccctaGGGAAGACTCCCCAGTGAC |
| PVX::gMYB21 | Fw: gaggtcagcaccagctagcaATTTTATAGTAAAGCAAGTTGTTTTAG<br>Rv: acatcaatagGGGAAGACTCCCCAGTGAC<br><br>Fw: gagtcttcccCTATTGATGTTATCATGGCAGTTTTAGAG<br>Rv: gggaaacttaacaaaccctaGGGAAGACTCCCCAGTGAC |
| PVX::gMYB24 | Fw: gaggtcagcaccagctagcaAATTCAGTCTTTAATCAGCCGTTTTAG<br>Rv: atactgctaatGGGAAGACTCCCCAGTGAC<br><br>Fw: gagtcttcccATTAGCAGTATATCGACTTAGTTTTAGAG<br>Rv: gggaaacttaacaaaccctaGGGAAGACTCCCCAGTGAC |

**Supplementary Table 2**: Primers employed for sgRNA2.1 construction and adaptation for PVX recombinant plasmids through Gibson assembly.

| Gene | Primer Sequence5'-3' |
|---|---|
| *NbDFR* | Fw: TTCATCTGCGCATCCCATCA<br>RV: TCCCTACTGAGTTTAAAGGTATCGA |
| *NbODO1* | Fw:AATTAAAACTGTCGTCCCAAG<br>Rv:GTCTGTCTCAGACCTTCCTGCTG |
| *NbMYB21* | Fw:TGCCAGGAAGAACAGATAACGAGA<br>Rv:ACGACATATGGCTACTGCTTCCT |
| *NbMYB24* | Fw:ATCATCAACAAGCTAGTACAAGC<br>Rv:TCATTTATTTCGTAAACAATTCATGGT |
| *NbF-box* | Fw:GGCACTCACAAACGTCTATTTC<br>Rv:ACCTGGGAGGCATCCTGCTTAT |

**Supplementary Table 3**: Primer pairs for RT-qPCR analyses.

| **Assay** | **Reference Gene (F-Box)** | **C- sample (TF measurement)** | **TF induced sample** |
|---|---|---|---|
| NbODO1 | 25,36 ± 0,58 | 31,23 ± 0,41 | 21,48 ± 1,79 |
| NbMYB21 | 25,94 ± 0,12 | 32,91 ± 0,70 | 25,40 ± 0,60 |
| NbMYB24 | 25,06 ± 0,23 | 34,88 ± 0,92 | 24,43 ± 0,22 |

**Supplementary Table 4**: Ct values obtained from the qRT-PCR assay for evaluating the transcriptional expression of *NbODO1*, *NbMYB21*, *NbMYB24* pPVX-gRNA induced samples.

| **Compound name** | **Fragmentation F2** |
|---|---|
| 2,4-Diisopropylphenyl acetate | 178, 163,117,147,105,91 |
| Methoxycalamenee | 217, 200, 157, 141 |
| Phenol 3,5 dimethyl | 122, 121, 107, 79 |
| 3,4,4-Trimethyl-2-cyclopenten-1-one | 109, 79, 81, 55, 70, 41 |
| Naphthalene, 1,2-dihydro-1,1,6-trimethyl | 172,163, 157,142, 115,128 |
| Naphthalene, 1,2-dihydro-4,5,7-trimethyl | 173, 157, 142 |
| α-Terpineol | 136, 121, 93, 81,59 |
| α-Terpinene | 136, 121, 93, 91, 77 |
| o-Cymene | 207, 119, 134, 91, 57 |
| α,4-Dimethyl-3-cyclohexene-1-acetaldehyde | 177, 135,91, 43 |

**Supplementary Table 5:** m/z fragmentation spectrum F2 of the differential metabolites obtained in the non-targeted analysis of the volatile content in negative control, *NbODO1, NbMYB21, NbMYB24* induced samples
